# Selective removal of visual working memory items at test

**DOI:** 10.1101/2025.05.22.655454

**Authors:** Chong Zhao, Temilade Adekoya, Sintra Horwitz, Edward Awh, Edward K. Vogel

## Abstract

Working memory (WM) tasks often require comparing remembered items to test displays, but little is known about how people selectively remove irrelevant information at test. Across three experiments, we used contralateral delay activity (CDA) to track WM load and examine selective removal. In Experiment 1, CDA amplitudes increased with set size even when only one item was probed, suggesting minimal removal based on spatial location. Experiment 2 ruled out spatial grouping by presenting items sequentially in the same location, yet more items were retained for larger set size. In Experiment 3, however, when items belonged to distinct mnemonic categories, CDA amplitudes at test were reduced, consistent with selective removal based on category relevance. Additionally, P3 old-new effects showed that decision speed and strength were influenced by the number of items maintained. Together, these results suggest that people selectively remove WM contents based on categorical relevance, not spatial cues, enabling more efficient memory-based decisions.

## Introduction

Visual working memory (VWM) is a limited-capacity system for temporarily representing task-relevant information so that it may be acted upon (Luck and Vogel 1997; Cowan 2001).While the encoding and storage mechanisms of VWM have been extensively examined and debated (Vogel et al. 2001; Alvarez and Cavanagh 2004; Xu and Chun 2006; Woodman and Vogel 2008; Zhang and Luck 2008; Harrison and Tong 2009), the mechanisms by which these VWM representations may be acted upon are less understood. That is, while information about multiple items can be concurrently maintained in VWM, it is clear that we can restrict our overt behavioral responses to selectively report just a single item from within the maintained set of items. Indeed, typical measures of VWM test memory by randomly choosing a single item from the remembered set and ask participants to decide whether this probed item is the same or different from its original value (Luck and Vogel 1997; Vogel and Machizawa 2004; Vogel et al. 2005). To do so, an individual must be able to restrict their behavioral response to just the one probed item in memory, a process that is referred to as *removal* (Lewis-Peacock et al. 2018; Oberauer 2018).

Theoretical models suggest that removing irrelevant items, particularly when individuals explicitly know those items are no longer needed, can enhance behavioral performance on working memory tasks (i.e., SOB-CS model, Oberauer et al., 2012). Empirically, Ecker, Oberauer, and Lewandowsky (2014) demonstrated this effect in a task where participants had to update the contents of their working memory with new letters. A red, bold cue signaled which information to update, appearing either briefly (200 ms) or after a longer delay (1500 ms) before the test. The key finding was that participants responded significantly more slowly when the cue-test interval was short compared to when it was long. This suggests that having more time allows participants to remove outdated information from memory, leading to faster and more efficient updating of the new memory array. Moreover, the benefit of removing irrelevant information extends beyond the immediate contents of working memory (Dames et al. 2025; Li, Frischkorn, Dames, et al. 2025; Li, Frischkorn, and Oberauer 2025). Oberauer and Lewandowsky (2016) found that when participants were exposed to a series of distractors before receiving the actual memory list, performance on the subsequent working memory task improved if there was a longer delay between the distractors and the memory list. Because the distractors occurred before the memory task began, these improvements are attributed to participants using the available free time to actively remove the distractors, thereby reducing their interference with later encoding. In the removal literature discussed above, distractors, or newly updated items, are typically presented at a different time than the target items; that is, targets and distractors are rarely shown simultaneously. An open question is whether people can selectively remove information from working memory even when the target and irrelevant items are presented within the same temporal window. Behavioral evidence supports this hypothesis. For instance, when participants were shown two items and later retro-cued to forget one, the precision of the remaining item in visual working memory was higher than when no such cue was given (Williams et al., 2012). Neural evidence further suggests that during the maintenance phase, when participants were instructed to forget certain items from the original memory array, their neural load was reduced. This indicates that they selectively removed some items from memory during maintenance, likely to enhance behavioral performance (Williams & Woodman, 2012).

However, while there appears to be consensus that some form of selective removal must occur, prior research has focused on the maintenance phase of visual working memory. Therefore, the specific mechanisms by which it is implemented at test are not well characterized. In particular, one plausible implementation of selective removal is that it is accomplished via editing the contents of VWM down to just the item relevant to the decision, while untested items are removed from active memory. While a VWM removal process would produce a straightforward means for achieving efficient working memory performance, there is currently little empirical evidence for this claim. Relatedly, it is still unclear how people accomplish selective removal in the decision process for making a memory-based judgement in VWM tasks. That is, when an individual is shown a single item probe at test and must decide whether that item is old or new, it is necessary to compare the probe item with the representation of the item at that location in VWM. During this comparison phase, it is unclear whether selective VWM removal is happening and if so, what information guides people to remove untested items. Importantly, prior work suggests that preserving the original configural context at test can enhance performance, even when only a single probe item is presented (Jiang et al., 2000), likely through relational grouping mechanisms (Jiang et al., 2004). These findings raise the possibility that participants benefit from maintaining relational structure rather than removing untested items during the test phase.

Here we address this question by examining multiple electrophysiological signatures of VWM load during both retention and test phases of a working memory task, enabling a precise measurement of whether selective VWM removal takes place at test, and if so, what the boundary conditions are that trigger this removal mechanism.

### Overview of experiments

In the present study, we aimed to investigate the extent of ***selective removal*** that occurred during *test phase* of visual working memory tasks. In Experiment 1, we tested whether selective removal at test could occur when participants were provided with a spatial retrieval cue in a single-probe change detection task. To further probe if simple spatial grouping affects the occurrence of editing, Experiment 2 employed a sequential working memory paradigm to test whether selective removal could occur even when they were no longer spatially grouped together in the same frame, as suggested by the relational coding findings. In Experiment 3, we explored whether selective removal at test could occur when items belonged to different categories (e.g., colors vs. shapes). In all of our experiments, we examined the neural signatures of visual working memory to test whether memory storage and decision making changed as a consequence of selective removal at test.

## Experiment 1

### Method

In Experiment 1, we tested if participants selectively remove their visual working memory load at test when they were shown a single test probe. We employed a single-probe change detection task in Experiment 1 (Luck and Vogel 1997) to ensure participants perceived the same number of items at test regardless of how many items were presented during the encoding phase. In this task, participants were presented with only one test probe, with a 50% chance of a change, regardless of whether the encoding set size was 2 or 4 (**Fig. 1**). If participants were able to effectively remove their working memory to retain only the item that shared the same spatial location as the test probe, then the CDA amplitude should not differ between set sizes. In contrast, if they did not engage in such removal, CDA amplitudes would remain higher for set size 4 compared to set size 2. Additionally, we examined the amplitude and latency of the P3 old-new effect to assess whether the strength and speed of decision-making changed with increasing set sizes, even when only a single probe was presented at test.

**Fig 1.**
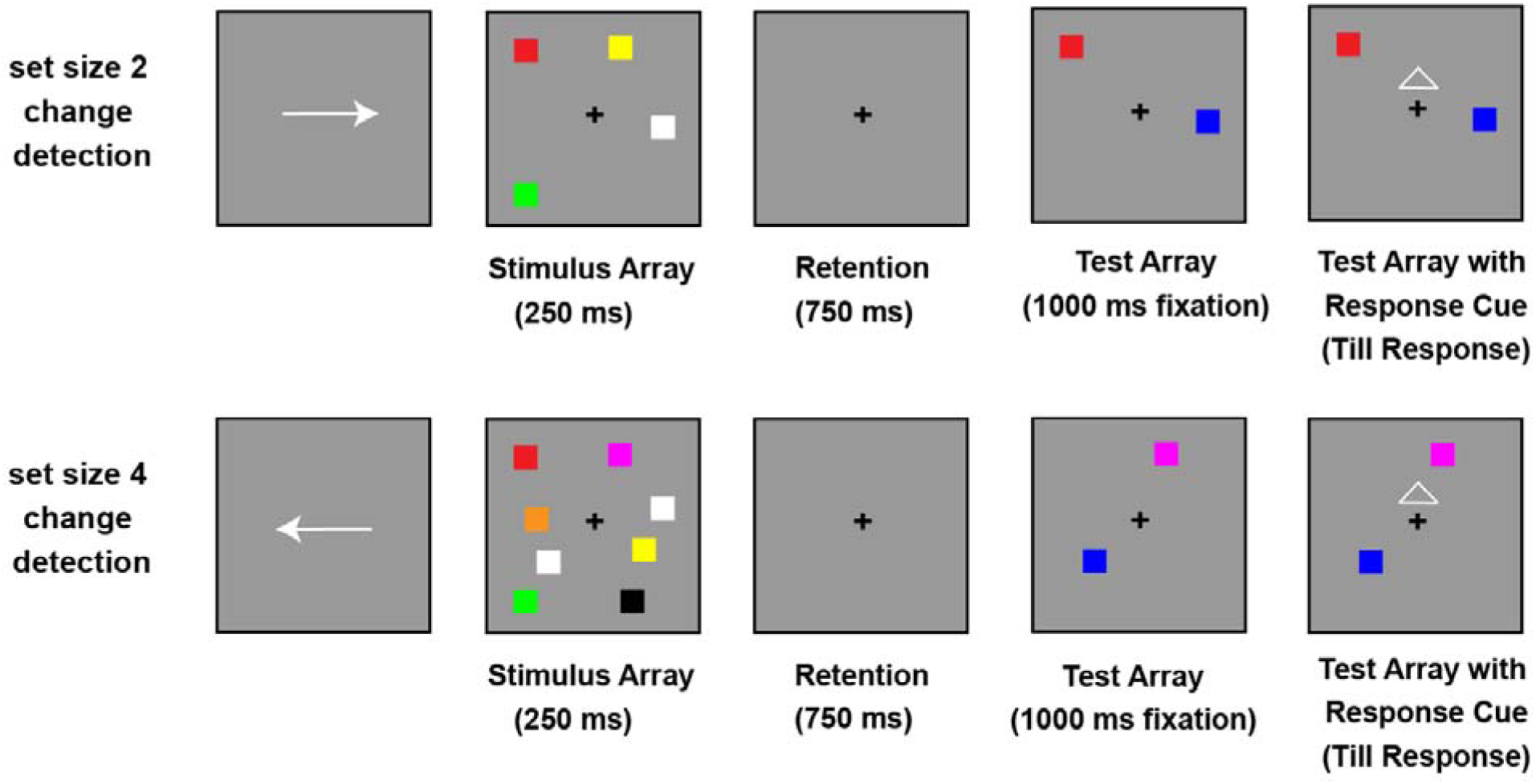
Change Detection Task (set size 2 and 4) used in Exp 1. Each trial began with a spatial cue (250 ms) indicating the relevant hemifield (left or right). This was followed by a memory array (250 ms) containing two or four colored squares per hemifield, drawn without replacement from a set of nine distinct colors. After a 750-ms delay, a test display appeared showing one colored square in each hemifield. The square on the cued side had a 50% chance of changing color, and participants were instructed to judge whether it had changed or not. The test display remained on screen until a response was made, but participants were required to maintain fixation and withhold all eye or motor movements for 1000 ms. Only after a movement cue appeared were they allowed to respond by pressing “z” (change) or “/” (no change) on the keyboard. Set size (2 or 4 items) was manipulated across trials.

### Participants

The experimental procedures were approved by the Institutional Review Board at The University of Chicago (IRB15-1290). All participants provided informed consent and were compensated with a cash payment of $20 per hour. They reported having normal color vision and normal or corrected-to-normal visual acuity. A total of 20 subjects were recruited, and no subject was rejected. This sample size was selected to ensure robust detection of set-size-dependent delay activity (Ngiam et al. 2021).

### Stimuli and Procedure

In Experiment 1, each trial started with a directional cue (250 ms) indicating which side of the screen (left or right) would be relevant. Following the cue, two or four colored squares (1.1° × 1.1°) briefly appeared (250 ms) in both hemifields, with at least 2.10° (1.5 objects) between them. The squares disappeared for a 750-ms delay. The squares were shown within a restricted area extending 3.1° horizontally from fixation and 3.5° vertically. The square colors were randomly selected from nine options (RGB values: red = 255, 0, 0; green = 0, 255, 0; blue = 0, 0, 255; yellow = 255, 255, 0; magenta = 255, 0, 255; cyan = 0, 255, 255; orange = 255, 128, 0; white = 255, 255, 255; black = 1, 1, 1). Colors were chosen without replacement within each hemifield, though colors could repeat across hemifields.

At test, a display containing one colored square in each hemifield appeared and remained on screen until a response was made. The square on the cued side had a 50% chance of changing its color, and the goal of the participants was to identify if the cued square had changed its color or not. The new color was always selected from the remaining colors in the set of nine possible colors that were not presented in the memory display. Participants were required to avoid making any eye or motor movements for 1000 ms after the test array was shown. Once a movement cue was presented, they pressed keys on the keyboard to indicate if the square had changed its color (“z”) or remained the same color as in the encoding array (“/”).

### Neural Index of VWM load (CDA)

Our intuitions and assumptions about selective removal have been strongly influenced by extensive work over the years examining the load of VWM during the maintenance period. There have been numerous demonstrations that individuals can voluntarily restrict access to VWM so that the most task relevant information may be represented in memory (Sperling 1960; Averbach and Coriell 1961; Vogel and Machizawa 2004; Vogel et al. 2005; Gazzaley and Nobre 2012). This work has provided compelling evidence that the selection during encoding and maintenance directly impacts what is stored in working memory. For example, when observers are presented with a display with equal numbers of objects in each hemifield, they can successfully restrict VWM storage to just the items in a spatially cued hemifield. This input gating is clearly apparent in an ERP component of human EEG known as the Contralateral Delay Activity (CDA), which shows a large sustained negative voltage over the hemisphere that is contralateral to the position of the to-be-remembered item as compared to the activity observed for the items in the ipsilateral hemisphere. Thus, the CDA reveals selective storage of just the task relevant items despite presence of irrelevant items in the uncued hemifield, indicating that observers can spatially gate the inputs to VWM when items are separated by hemifield. Furthermore, the amplitude of the CDA is sensitive to the number of task-relevant items stored in VWM within the cued hemifield. It rises in amplitude as a function of number of items but reaches an asymptote for arrays of 3-4 items, respecting both group and individual capacity limits (Cowan 2001; Vogel and Machizawa 2004; Collins and Frank 2012). Although CDA has been widely used metric of the current load in VWM research, prior studies have focused on the retention interval preceding the test, leaving unclear how many items are retained in VWM at test.

### Neural index of memory-based decisions (P3)

In addition to a reliable measure of VWM load (i.e., CDA), to examine the role of selective VWM removal in the decision process, it is useful to have a direct index of the latency and strength of the resulting decision. A key neural marker of this decision process is the P3 component (Donchin and Coles 1988), which is flexibly sensitive to task goals in recognition memory tasks. During these recognition tasks, the P3 shows a larger positive amplitude for old items compared to new items, a phenomenon referred to as the P3 old/new effect (Wilding and Rugg 1996; Friedman and Johnson Jr 2000; Rugg and Curran 2007; Vilberg and Rugg 2008). Moreover, the amplitude of the old/new difference has been repeatedly shown to be sensitive to the accuracy and confidence of the memory-based decision (Yonelinas 2001; Boldini et al. 2004; Curran 2004; Woodruff et al. 2006). Thus, the amplitude of this P3 old-new effect may serve as an index of the strength of working memory decisions in our visual working memory tasks. Furthermore, prior work has demonstrated that the *latency* of the P3 component increases with VWM set size, suggesting that larger set sizes impose greater demands on decision processes (Hyun et al. 2009). The latency of P3 component reflected how fast participants accumulated evidence towards a motor response and had been shown to be predictive of response time measures (Roth et al. 1978; Giedke et al. 1981; Verleger et al. 1991; Verleger 1997; Doucet and Stelmack 1999). Thus, the P3 provides an explicit index of when decisions are made, enabling direct conclusions about whether VWM removal occurs prior to decision making.

### CDA and P3 Analysis

In Experiment 1, event-related potentials were computed using a 250-ms pre-stimulus baseline. Averaged activity at each electrode was used to calculate contralateral and ipsilateral amplitudes for predefined electrode pairs based on previous research: O1/O2, PO3/PO4, PO7/PO8, P3/P4, and P7/P8 (Ngiam et al. 2021). Data were analyzed without additional filtering beyond the initial .01-30 Hz online filter.

To compute the contralateral delay activity (CDA), we examined the voltage difference between parieto-occipital electrodes on the side opposite to the cued direction and those on the same side. During the retention phase, we focused on the 500ms to 1000ms time window after the onset of the encoding array, a period when only the fixation cross was displayed on the screen. For the comparison phase, our window of interest was from 1500ms to 2000ms after the onset of the encoding array, when the test array appeared, but participants were instructed to refrain from making any eye movements or motor responses.

The amplitude of the P3 old/new component was measured at electrodes P3, P4, P7, P8, and Pz, selected based on prior research (Rugg & Curran, 2007). We computed the average ERP separately for old and new trials and then derived the P3 old/new effect as the difference wave (old minus new) for each set size. The analysis focused on the comparison phase, with a time window of interest from 1250 to 1550 ms, corresponding to 250–550 ms after test array onset, as informed by previous literature (Hyun et al., 2009). Amplitude results were aggregated across the time window of interest and statistical significance were reported based on the mean amplitudes of the P3 old/new effect for each participant between different set sizes. To estimate P3 old/new latency, we employed a jackknife-based approach using the 50% area under the curve method, reporting the mean latency across jackknife subsamples for each condition. We reported both the amplitude and latency of the P3 old/new effect. As noted by Luck (2014), these two measures are interdependent: changes in amplitude can reflect underlying shifts in latency because ERPs represent averages across single trials. Accordingly, we caution against interpreting amplitude and latency entirely independently. By presenting both measures alongside this caveat, we aim to provide a more complete characterization of the memory-related decision-making processes captured by the P3 old/new effect.

### Result

Behaviorally, we observed a clear set size effect in accuracy. Performance was lower for set size 4 (mean accuracy = 78.2%, mean RT = 1109 ms) than for set size 2 (mean accuracy = 89.7%, mean RT = 1007 ms, t(19) = 7.01, p < 0.01, for detailed RT by old/new trials see **Supp Table 1**).

During the retention phase, we hypothesized that participants would hold all items from the array within the limits of their working memory capacity (Luck and Vogel 1997). Consequently, we expected CDA amplitudes in the set size 2 condition to be significantly lower than in the set size 4 condition. Consistent with previous results from the change detection paradigm, we observed a significant set size effect during the retention interval (*t*(19) = 5.24, *p* < 0.01, see **Fig. 3**), where the CDA amplitude for set size 2 was significantly lower than for set size 4. This suggests that participants retained more items at higher set sizes (set size 4 > set size 2)from the array throughout the retention period.

When the test array was presented, if participants were selectively accessing only the changed item (i.e., *spatially-guided removal* of irrelevant items in their working memory), we would expect no significant difference in CDA amplitudes between the set sizes. However, contrary to the *spatially-guided removal* hypothesis (see **Fig. 2**), we found a significant difference in CDA amplitudes between set size 2 and set size 4 arrays during the comparison phase. Specifically, CDA amplitudes were significantly higher for set size 4 compared to set size 2 (*t*(19) = 4.00, *p* < 0.01, see **Fig. 3**), indicating that participants retained more items for comparison in the larger arrays. A two-way ANOVA (phase x set size) revealed that CDA amplitudes were significantly higher for set size 4 than set size 2 (F(1,19) = 20.53, *p* < 0.01), and higher for retention phase than comparison phase (F(1,129 = 27.32, *p* < 0.01), and an interaction between these two factors (F(1,19) = 14.31, *p* < 0.01). Collectively, these findings suggested that untested items remained during the comparison phase, and participants did not remove all spatially irrelevant items and keep the single working memory item at test corresponding to the test probe’s spatial location.

**Fig 2.**
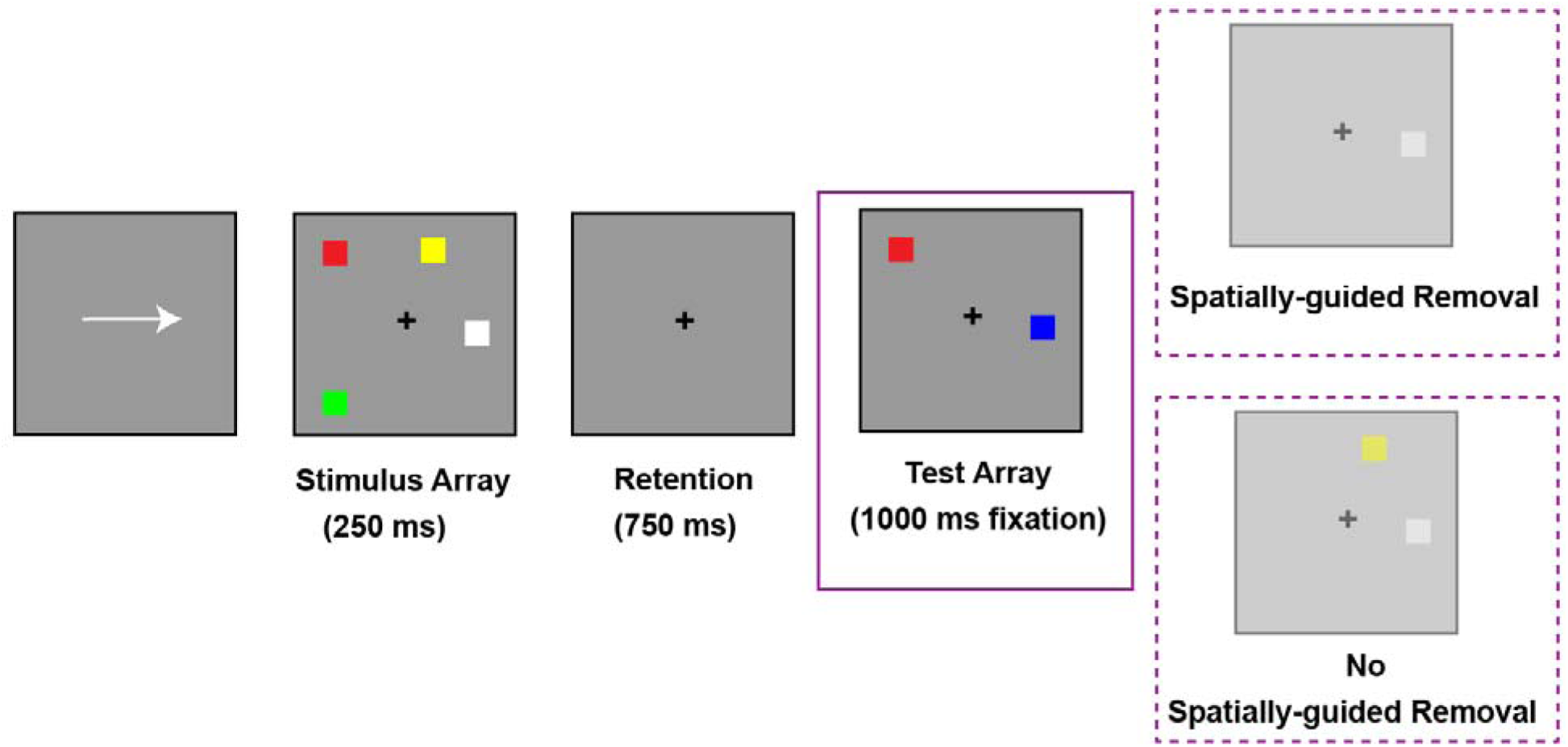
Spatially-guided Removal vs No Removal hypotheses at test in Exp 1. If participants actively engaged in spatially guided removal their visual working memory during the test phase, they should retain only a single item that shared the same location with the test probe by that point (upper). In contrast, if no such removal occurred, neural activity during the test phase should continue to reflect the number of items originally stored (lower).

**Fig 3.**
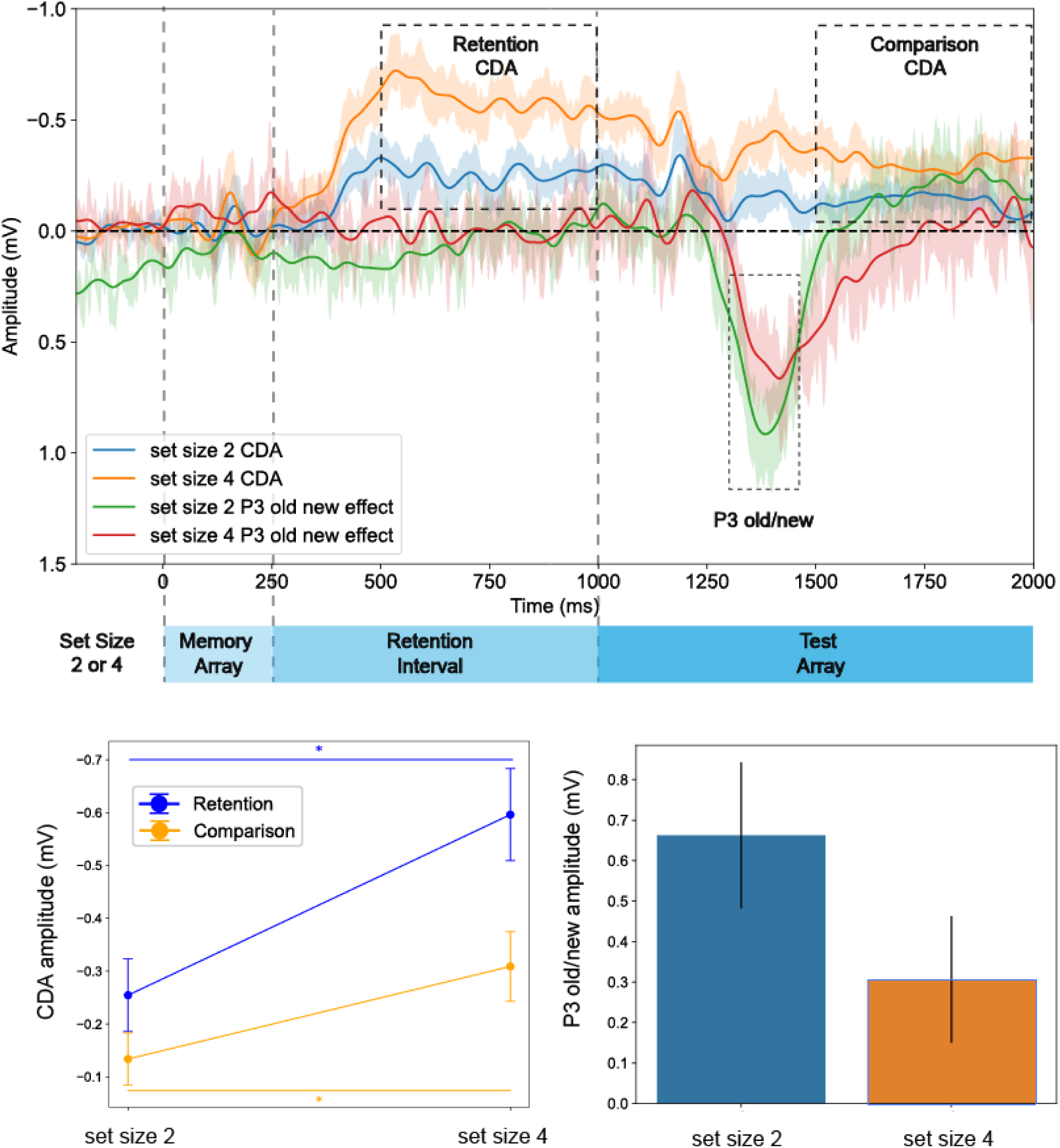
Retention and Comparison Phase CDA, and Comparison Phase P3 old new effect Results for Exp 1. CDA amplitude was larger for set size 4 compared to set size 2, suggesting that participants did not remove their working memory at test even when a single test probe was shown. Additionally, the test display evoked a larger and earlier P3 component for set size 2, indicating that participants made a faster and more robust decision in the set size 2 condition than in the set size 4 condition. These results suggest that participants did not engage in active removal of visual working memory contents based on spatial locations at test.

To further examine whether participants removed the items in visual working memory at test for decision-making, we compared the amplitude of the P3 old-new effect between set sizes 2 and 4. Between 250 and 450 ms after test array onset, the P3 old-new effect was larger for set size 2 than for set size 4 (*t*(19) = 2.20, *p* = 0.04), suggesting that decisions were more robust when fewer items were maintained in working memory. Additionally, P3 latencies were significantly faster for set size 2 (395.06 ms) than for set size 4 (451.09 ms, p < 0.01). These findings indicate that decision-making was slower for larger encoding set sizes, suggesting that removal of all irrelevant items that did not share the same location with the test probe failed to occur at test, even though participants were able to make accurate decisions.

## Experiment 2

### Method

Our Experiment 1 suggested that people did not remove their working memory based on spatial relevance even when only a single test probe was shown at test. One possible reason participants were keeping more than one item is that arrays with larger set sizes include more spatial locations on the screen due to spatial grouping effect (i.e., Musfeld et al., 2024). In a change detection setting, participants used relational coding to encode neighboring objects as part of an ensemble (Bateman, 2018). As a result, participants may access a larger ensemble for set size 4 compared to set size 2 when performing the working memory comparison. To test whether spatial grouping contributed to the failure of *spatially-guided removal*, we introduced a third condition in Experiment 2, using a set size 4 in a sequential working memory paradigm (Vogel et al. 2005). In this design, participants first encoded one set size 2 array, followed by a second set size 2 array, with both arrays presented at the same spatial locations. This meant that each spatial position was associated with two different colored squares. If spatial grouping was driving the lack of *spatially-guided removal* effect, we would expect to observe a CDA equivalent to set size 2 for this sequential set size 4 condition, as only two locations were memorized. However, if the CDA is similar to that of the standard set size 4 condition, we could rule out a simple spatial grouping strategy as the factor influencing the fate of untested working memory items during the comparison phase.

### Participants

The experimental procedures were approved by the Institutional Review Board at The University of Chicago (IRB15-1290). All participants provided informed consent and were compensated with a cash payment of $20 per hour. They reported having normal color vision and normal or corrected-to-normal visual acuity. A total of 20 subjects were recruited, and 1 subject was rejected due to excessive eye movements. This sample size was selected to ensure robust detection of set-size-dependent delay activity (Ngiam et al. 2021).

### Stimuli and Procedure

In Experiment 2, each trial started with a directional cue (250 ms) indicating which side of the screen (left or right) would be relevant (see **Fig. 4**). For the set size 2 and 4 conditions, the procedure was the same as in Experiment 1. For the sequential set size 4 condition, two colored squares (1.1° × 1.1°) briefly appeared (250 ms) in both hemifields, with at least 2.10° (1.5 objects) between them. Followed by a 250-ms retention interval, two more colored squares briefly appeared (250 ms) in the same spatial locations as the first encoding array, and the participants were instructed to memorize all four colors in these two arrays. The colors used by the second array were different from the two used in the first array, resulting in four unique colors to be remembered. The squares then disappeared for a 250-ms delay. The squares were shown within a restricted area extending 3.1° horizontally from fixation and 3.5° vertically. The square colors were randomly selected from nine options (RGB values: red = 255, 0, 0; green = 0, 255, 0; blue = 0, 0, 255; yellow = 255, 255, 0; magenta = 255, 0, 255; cyan = 0, 255, 255; orange = 255, 128, 0; white = 255, 255, 255; black = 1, 1, 1). Colors were chosen without replacement within each hemifield, though colors could repeat across hemifields.

**Fig 4.**
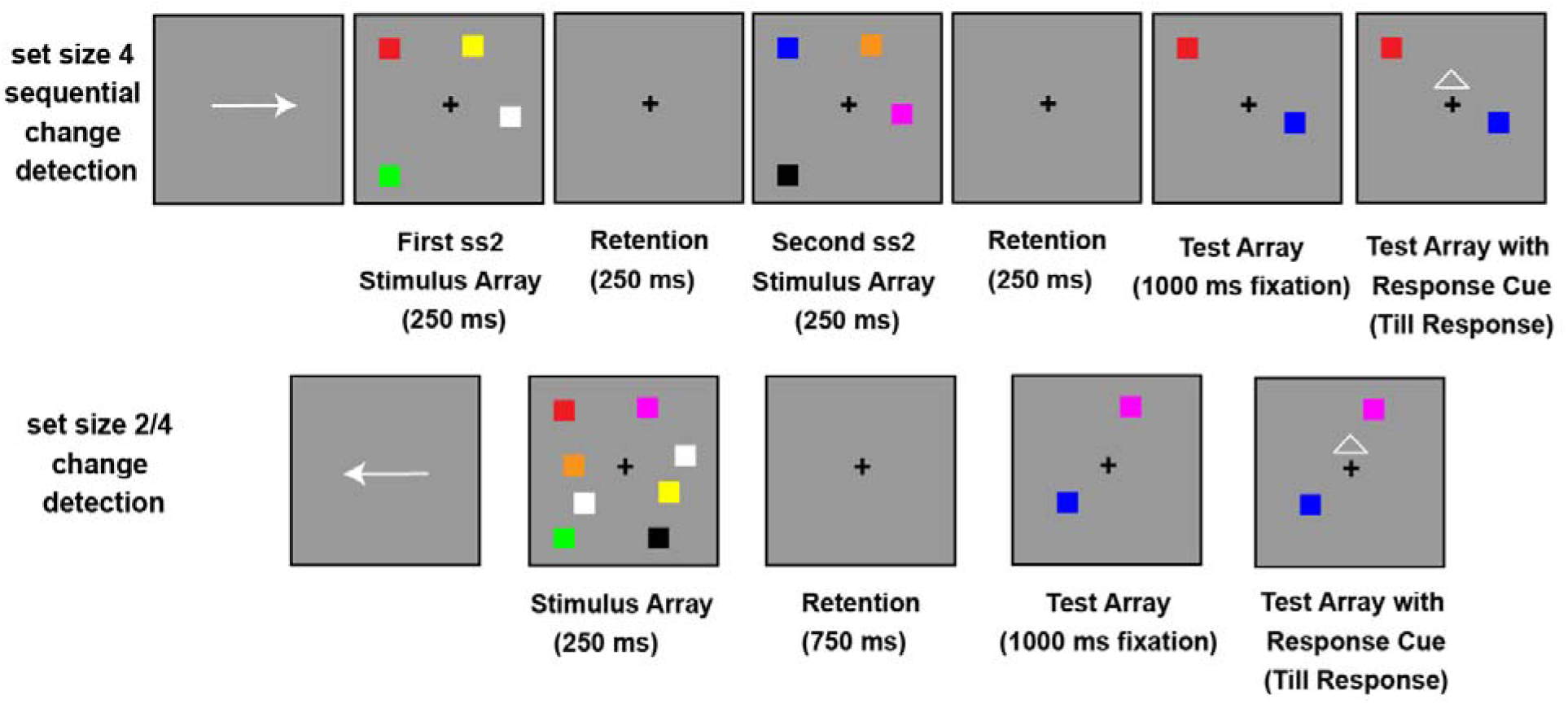
Change Detection Task (set size 2 and 4) used in Exp 2. Each trial began with a spatial cue (250 ms) indicating the relevant hemifield (left or right). This was followed by a memory array (250 ms) containing either two or four colored squares per hemifield, drawn without replacement from a set of nine distinct colors. In the sequential condition, the four-item memory array was split into two sequential displays of two items each, separated by a brief interstimulus interval (250 ms), creating a set size 2 + set size 2 condition. After a 750-ms delay (250 ms after the second array for the sequential condition), a test display appeared showing one colored square in each hemifield. The square on the cued side had a 50% chance of changing color, and participants were instructed to judge whether it had changed. The test display remained on screen until a response was made, but participants were required to maintain fixation and withhold all eye or motor movements for 1000 ms. Only after a movement cue appeared were they allowed to respond by pressing “z” (change) or “/” (no change) on the keyboard. Set size (2, 4, or 2+2 items) was manipulated across trials.

At test, a display containing one colored square in each hemifield appeared and remained on screen until a response was made. The square on the cued side had a 50% chance of changing its color, and the goal of the participants was to identify if the cued square had changed its color or not. For the sequential set size 4 conditions, a changed color would be different from both colored squares that were placed at the same location as the test probe. Similarly, in the same color trials, the color of the test probe would be the same as one of the two colored squares in the encoding array that shared the same spatial location with the test probe. Participants were required to avoid making any eye or motor movements for 1000 ms after the test array was shown. Once a movement cue was presented, they pressed keys on the keyboard to indicate if the square had changed its color (“z”) or remained the same color as in the encoding array (“/”).

### CDA Analysis

In Experiment 2, For the set size 2 and 4 conditions, the analyses were identical to Experiment 1. For the sequential set size 4 condition, we focused on 500ms to 750ms for the first encoding array of set size 2, and 750ms to 1000ms for the second encoding array of set size 2. Across all conditions, our window of interest was from 1500ms to 2000ms after the onset of the encoding array for the comparison phase, when the test array appeared, but participants were instructed to refrain from making any eye movements or motor responses.

## Result

Behaviorally, we observed a clear set size effect in accuracy. Performance was lower for both set size 4 (mean accuracy = 79.2%, mean RT = 778 ms, t(18) = 16.00, p < 0.01) and sequential set size 4 conditions (mean accuracy = 80.8%, mean RT = 740 ms, t(18) = 9.12, p < 0.01) than for set size 2 (mean accuracy = 91.4%, mean RT = 718 ms, for detailed RT by old/new trials see **Supp Table 2**). Additionally, the accuracies in sequential set size 4 condition were not different from those in set size 4 condition (t(18) = 1.31, p = 0.21).

During the retention phase, we hypothesized that participants would hold all items from the array within the limits of their working memory capacity (Luck and Vogel 1997). Similar to Experiment 1, we expected CDA amplitudes in the set size 2 condition to be significantly lower than in the set size 4 condition. Replicating our previous findings, we observed a significant set size effect during the retention interval (*t*(18) = 4.13, *p* < 0.01), where the CDA amplitude for set size 2 was significantly lower than for set size 4. This suggests that participants retained more items at higher set sizes (set size 4 > set size 2) from the array throughout the retention period. Similar to the sequential change detection paradigm in previous studies (Vogel et al. 2005), we found that the CDA for the first set size 2 array was similar to the set size 2 CDA (*t*(18) = 0.11, *p* = 0.91, see **Fig. 5**), suggesting that participants first encoded all two items into their working memory during the sequential encoding. After the onset of the second set size 2 array, we found that the resulting CDA for the sequential condition was similar to the set size 4 condition (*t*(18) = 0.66, *p* = 0.52, see **Fig. 5**) and way higher than the set size 2 condition (*t*(18) = 2.87, *p* < 0.01, see **Fig. 5**). These suggested that although the two arrays overlapped in spatial locations, participants were able to encode all 4 items into their working memory, resulting in a CDA similar to the simultaneous set size 4 condition.

**Fig 5.**
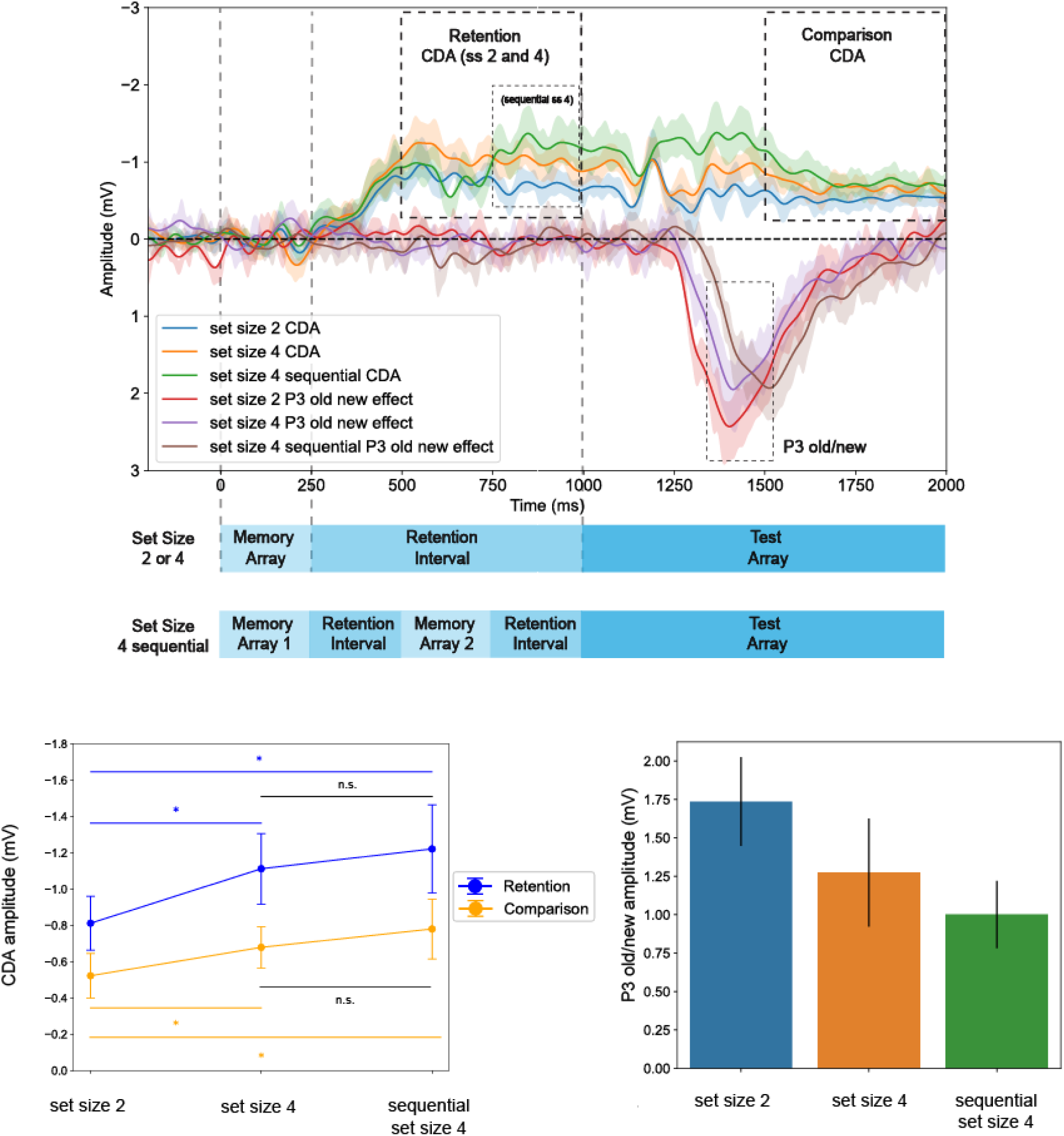
Retention and Comparison Phase CDA Results, and Comparison Phase P3 old new effect for Exp 2. CDA amplitude was larger for set size 4 compared to set size 2, consistent with the maintenance of more items in visual working memory at test. Notably, the sequential condition (set size 2 + 2) produced CDA amplitudes comparable to the simultaneous set size 4 condition, suggesting that participants kept 4 items in working memory even if they were encoded in two different temporal frames in the sequential condition. Additionally, the test display evoked a larger and earlier P3 component for set size 2 condition than the sequential and simultaneous set size 4 conditions, suggesting that people did not remove their working memory at test even across distinct temporal frames in the sequential condition. These results suggest that participants did not engage in active removal of their visual working memory contents based on spatial relevance at test.

When the test array was presented, if participants performed *spatially-guided removal* down to the one encoded item that shared the same spatial location as test probe, we would expect no significant difference in CDA amplitudes between the set sizes. Refuting this hypothesis and replicating our findings in Experiment 1, we found a significant difference in CDA amplitudes between set size 2 and set size 4 arrays during the comparison phase. Specifically, CDA amplitudes were significantly higher for set size 4 compared to set size 2 (*t*(18) = 3.97, *p* < 0.01, FDR-corrected *q* < .05, see **Fig. 5**), indicating that participants retained more items for comparison in the larger arrays. To further test if participants were able to selectively retrieve the set of items that shared the spatial positions with the test probe, we examined the CDA for the sequential array during the comparison phase. If participants were able to select the items that shared the same positions with the test probe, the resulting CDA would be similar to set size 2 condition. Contrary to this *spatially-guided removal* hypothesis, we showed that the CDA for sequential set size 4 condition was the same as simultaneous set size 4 condition (*t*(18) = 0.52, *p* = 0.61), and significantly higher than set size 2 condition (*t*(18) = 2.31, *p* = 0.03, FDR-corrected *q* < .05, see **Fig. 5**). Our findings further challenged the *spatially-guided removal* hypothesis in that participants seemed to retrieve more items if they encoded them at first, regardless of whether overlapping spatial or temporal markers were used for the items. Therefore, untested items remained in visual working memory during the comparison phase. Furthermore, simple spatial grouping strategy did not explain the lingering of untested items we observed in Experiments 1-2. We concluded that people did not remove their working memory load based on spatial relevance at.

To further examine whether participants remove their visual working memory contents based on spatial relevance for decision-making, we compared the amplitude of the P3 old-new effect between set sizes 2, 4 and sequential 4. Between 250 and 550 ms after test array onset, the P3 old-new effect was larger for set size 2 than for set size 4 (*t*(18) = 2.11, *p* = 0.048), suggesting that decisions were more robust when fewer items were maintained in working memory. Similarly, the P3 old-new effect was larger for set size 2 than for sequential set size 4 (*t*(18) = 2.23, *p* = 0.04), indicating weaker evidence for decision making for set size 4 regardless of simultaneous or sequential encoding. Lastly, set size 4 and sequential set size 4 did not differ in P3 old-new amplitude (*t*(18) = 0.82, *p* = 0.42).

In addition to the P3 old-new amplitude, we replicated our findings in Experiment 1 that P3 latencies were significantly faster for set size 2 (443.26 ms) than for set size 4 (461.68 ms, *p* < 0.01). Moreover, we found that sequential set size 4 had a significantly slower latency than either set size 2 or 4, suggesting that the participants spent more time accessing information from the second array (528.84ms, *p* < 0.01). These findings replicated our findings in Experiment 1 that decision-making was slower for larger encoding set sizes, and *spatially-guided removal* was not playing a significant role in selecting the one spatially relevant item.

## Experiment 3

### Method

Experiments 1 and 2 showed that people did not *remove* their working memory contents based on spatial relevance during test phase with color-only encoding arrays. Furthermore, Experiment 2 suggested that simple spatial grouping was not the reason participants consistently kept more than one item at test (i.e., fail to accomplish *spatial removal*). However, it is plausible that participants can remove certain working memory items at test on the basis of some other rule, such as stimulus category. Indeed, previous work has shown a small behavioral advantage for working memory tasks with a mixed set of categories over homogeneous displays of all colors (Cai et al. 2020; Wennberg and Serences 2024). This advantage could be driven by more efficient removal in which the items from the untested category can be edited to reduce potential impact on the decision. Though, it is similarly plausible that this advantage could be the result of increased storage for heterogeneous arrays over homogeneous arrays (Cohen et al. 2014). If this were the case, we would expect to see an increased CDA amplitude during the retention period for mixed category displays over homogenous displays of the same set size.

Experiment 3 used a mixed array design in which participants encoded two colors and two shapes during a mixed set size 4 condition (see **Fig. 6**). The test probe in each trial was either a colored square or a shape, and participants were informed that colors would only change to other colors, and shapes would only change to other shapes. If mnemonic context influenced the fate of the untested items, participants would employ *category-based removal* strategy in restricting access to the subset of two stimuli in the array relevant to the category of the test probe, resulting in a CDA equal to set size 2. Conversely, if participants were unable to filter items by mnemonic context (i.e., use *category-based removal*), we would observe a CDA equal to set size 4 in the mixed array condition, indicating that the entire encoding array, regardless of tested or untested, remained in working memory during test (see **Fig. 7**).

**Fig 6.**
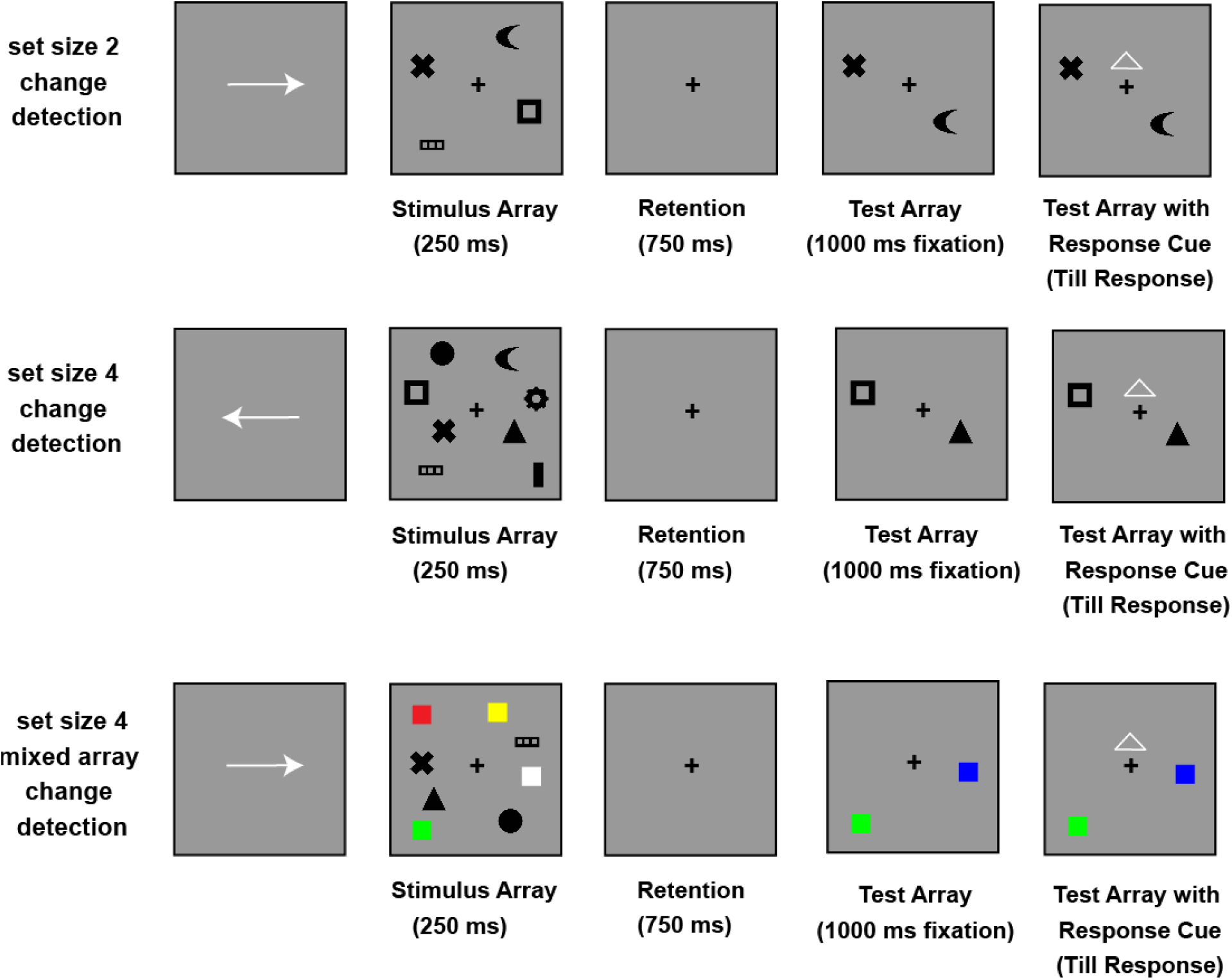
Change Detection Task (set size 2, 4 and mixed set size 4) used in Exp 3. Each trial began with a directional cue (250 ms) indicating the relevant hemifield (left or right). This was followed by a memory array (250 ms) containing either two shapes (set size 2), four shapes (set size 4), or a mixed array of two colored squares and two shapes (set size 4 mixed). After a 750-ms delay, a test display appeared with one item per hemifield. In the shape-only conditions, a shape appeared at test; in the mixed condition, either a color square or a shape appeared, depending on the item originally in that location. The test item on the cued side had a 50% chance of changing its feature (color or shape). Participants were instructed to judge whether the test probe matched the item previously shown at the same location. They were asked to maintain fixation and refrain from making eye or motor movements for 1000 ms after the test array onset. Once a movement cue appeared, they responded using the keyboard: “z” for a change, and “/” for no change.

**Fig 7.**
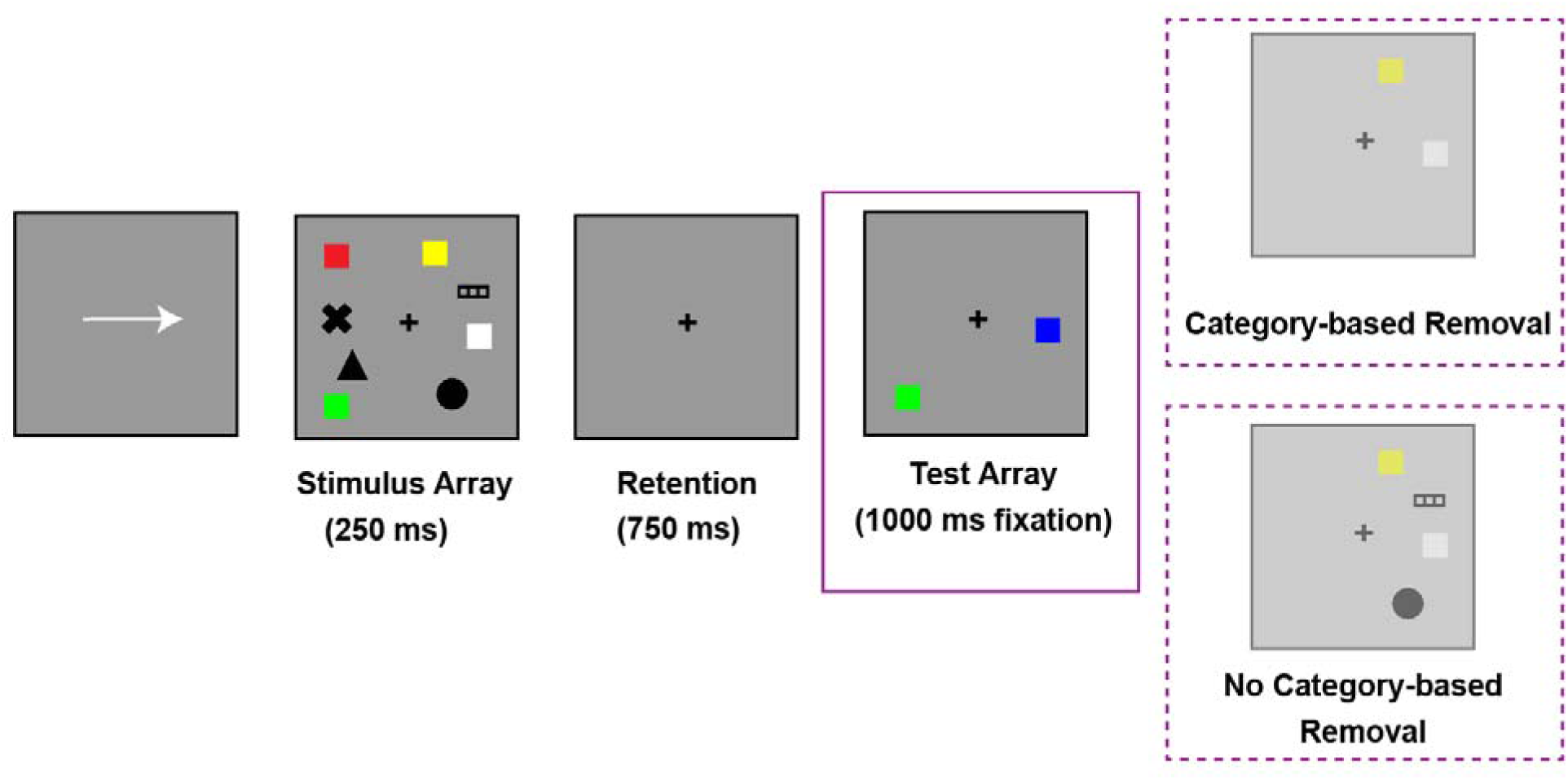
Categorical-based Removal vs No Categorical-based Removal hypotheses in Exp 3. If participants actively remove irrelevant items based on category during the test phase, they should only retain items that shared the same category as the test probe (upper). In contrast, if no category-based removal occurred, neural activity during the test phase should continue to reflect the number of all items encoded regardless of categorical relevance (lower).

### Participants

The experimental procedures were approved by the Institutional Review Board at The University of Chicago (IRB15-1290). All participants provided informed consent and were compensated with a cash payment of $20 per hour. They reported having normal color vision and normal or corrected-to-normal visual acuity. A total of 25 subjects were recruited, and 1 subject was rejected due to excessive blocking and noise trials. This sample size was selected to ensure robust detection of set-size-dependent delay activity (Ngiam et al. 2021).

### Stimuli and Procedure

In Experiment 3, each trial started with a directional cue (250 ms) indicating which side of the screen (left or right) would be relevant. Following the cue, two shapes (set size 2), four shapes (set size 4), or two color squares and two shapes (set size 4 mixed array, 1.1° × 1.1°) briefly appeared (250 ms) in both hemifields, with at least 2.10° (1.5 objects) between them. The encoding array disappeared for a 750-ms delay. Stimuli in the encoding array were shown within a restricted area extending 3.1° horizontally from fixation and 3.5° vertically. For the color squares, the colors were randomly selected from nine options (RGB values: red = 255, 0, 0; green = 0, 255, 0; blue = 0, 0, 255; yellow = 255, 255, 0; magenta = 255, 0, 255; cyan = 0, 255, 255; orange = 255, 128, 0; white = 255, 255, 255; black = 1, 1, 1). For the shapes, the shapes were randomly selected from nine options (Zhao et al. 2022). Shapes and colors were chosen without replacement within each hemifield, though they could repeat across hemifields.

At test, a display containing one stimulus (shape in the set size 2 and 4 condition, color or shape in the set size 4 mixed array condition) in each hemifield appeared and remained on screen until a response was made. The stimulus on the cued side had a 50% chance of changing its feature value (either color or shape), and the goal of the participants was to identify if the test probe had changed its feature value (color or shape) compared to the item in the same location during encoding. Participants were required to avoid making any eye or motor movements for 1000 ms after the test array was shown. Once a movement cue was presented, they pressed keys on the keyboard to indicate if the test probe had changed its color/shape (“z”) or remained the same color/shape as in the encoding array (“/”).

### CDA Analysis

CDA analyses in Experiment 3 followed the same time window and electrodes as in Experiment 1.

## Result

Behaviorally, we observed a clear set size effect in accuracy. Performance was lower for both uniform set size 4 (mean accuracy = 78.8%, mean RT = 629 ms, t(23) = 19.33, p < 0.01) and mixed set size 4 conditions (mean accuracy = 82.9%, mean RT = 604 ms, t(23) = 12.32, p < 0.01) than for uniform set size 2 (mean accuracy = 93.6%, mean RT = 524 ms, for detailed RT by old/new trials see **Supp Table 3**). Importantly, the mixed set size 4 condition showed a modest but reliable improvement relative to the uniform set size 4 condition (t(23) = 6.72, p < 0.01), replicating previous findings (Cai et al. 2020; Wennberg and Serences 2024).

During the retention phase, we hypothesized that participants would hold all items from the array within the limits of their working memory capacity (Luck and Vogel 1997). Similar to Experiments 1 and 2, we expected CDA amplitudes in the set size 2 condition to be significantly lower than in the set size 4 condition. Replicating our previous findings, we observed a significant set size effect during the retention interval (*t*(23) = 4.67, *p* < 0.01, see **Fig. 8**), where the CDA amplitude for set size 2 was significantly lower than for set size 4. This suggests that participants retained more items at higher set sizes (set size 4 > set size 2) from the array throughout the retention period. In our mixed array set size 4 condition, we found that the CDA for the mixed set size 4 array was higher than the set size 2 CDA (*t*(23) = 3.61, *p* < 0.01, see **Fig. 8**) and similar to the uniform set size 4 array (*t*(23) = 0.63, *p* = 0.54, see **Fig. 8**). These suggested that participants were able to encode all 4 items into their working memory for the mixed array, resulting in a CDA similar to the uniform set size 4 condition.

**Fig 8.**
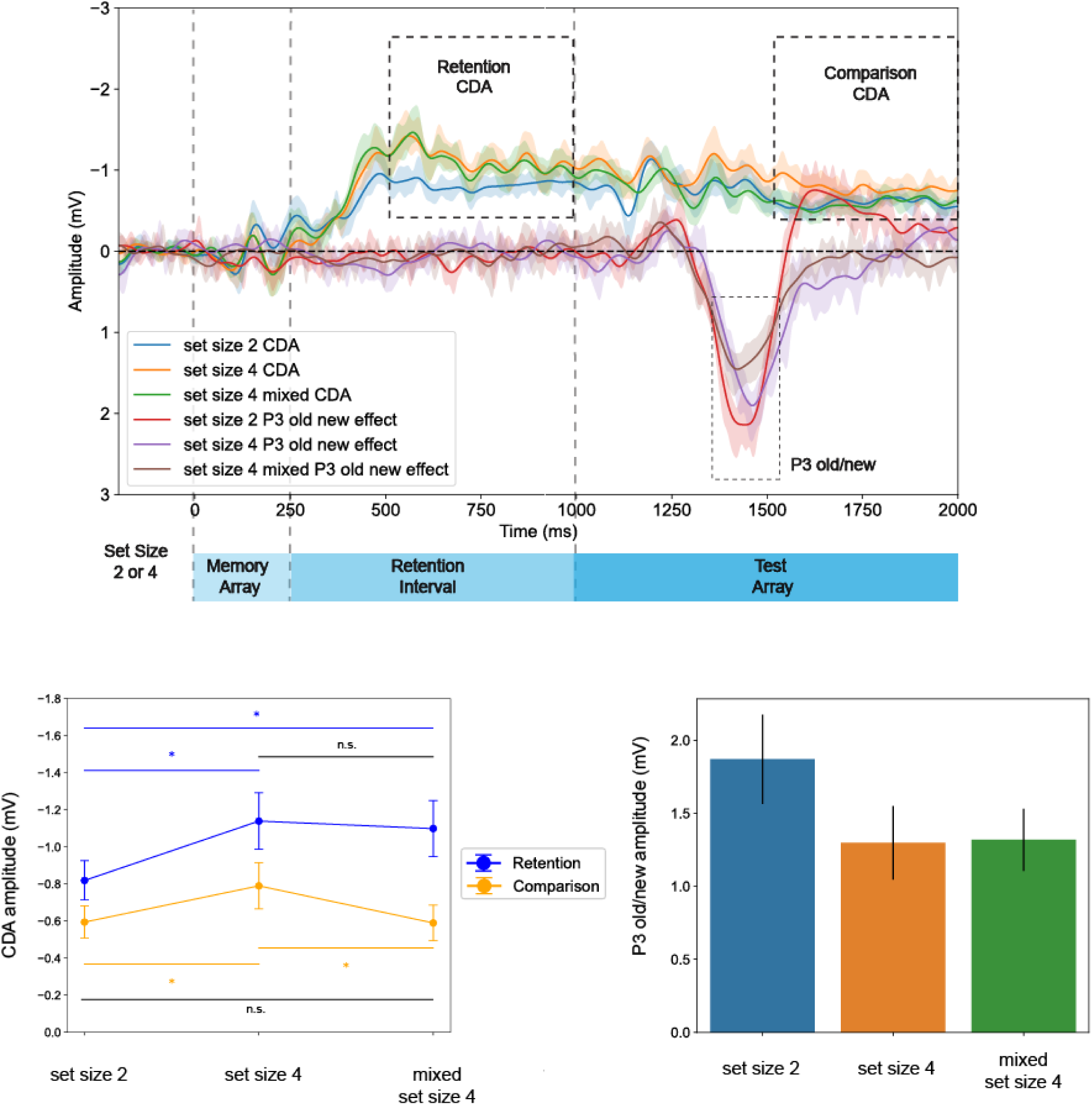
Retention and Comparison Phase CDA, and Comparison Phase P3 old new effect Results for Exp 3. CDA amplitude was larger for set size 4 compared to set size 2, consistent with the maintenance of more items in visual working memory at test. However, the mixed set size 4 condition produced CDA amplitudes comparable to the uniform set size 2 condition, suggesting that only two categorically relevant items were kept in working memory at test. Additionally, the test display evoked an earlier P3 component for mixed set size 4 condition than the uniform set size 4 conditions, suggesting that participants were making decisions based on two items instead of all four items at test. These results suggest that participants engaged in category-based removal of their visual working memory contents at test.

When the test array was presented, replicating our findings in Experiments 1 and 2, we found a significant difference in CDA amplitudes between set size 2 and set size 4 arrays during the comparison phase. Specifically, CDA amplitudes were significantly higher for set size 4 compared to set size 2 (*t*(23) = 2.31, *p* = 0.03, FDR-corrected *q* < .05, see **Fig. 8**), indicating that participants retained more items for comparison in the larger arrays.

In Experiment 3, we further aimed at testing if participants did not perform removal at all (i.e., no category-based *removal*), or instead actively remove the items that were mnemonically irrelevant to the test probe (i.e., category-based *removal*). If participants always had to access all items, we would expect that the CDA for mixed array set size 4 remained the same as uniform set size 4 array, even though two of the items belonged to a category that would never be relevant to the test probe. Alternatively, if participants were able to selectively remove the subset of items irrelevant to the category of the test probe, we would observe a lower CDA for mixed array set size 4 than uniform set size 4, and similar to the CDA amplitude in uniform set size 2 condition. To test if participants were able to selectively remove the subset of items that did not share the mnemonic context with the test probe, we examined the CDA for the mixed array during the comparison phase. We showed that the CDA amplitude in mixed set size 4 condition at test was significantly lower than in uniform set size 4 condition at test (*t*(23) = 2.95, *p* < 0.01, FDR-corrected *q* < .05, see **Fig. 8**), and similar to the CDA in uniform set size 2 condition during comparison (*t*(23) = -0.06, *p* = 0.95, see **Fig. 8**). These findings suggested that participants only selectively remove the untested items if they belonged to a different category from the test probe. Therefore, we concluded that participants actively performed category-based *removal* at test when they realized that only items from the relevant category were useful for the working memory test.

To further examine whether this category-based *removal* of items affects decision-making, we compared the amplitude of the P3 old-new effect between set sizes 2, 4 and mixed set size 4. Between 350 and 450 ms after test array onset, the P3 old-new effect was larger for set size 2 than for set size 4 (*t*(23) = 2.59, *p* = 0.02), suggesting that decisions were more robust when fewer items were maintained in working memory. Similarly, the P3 old-new effect was larger for set size 2 than for mixed set size 4 (*t*(23) = 2.20, *p* = 0.04), indicating weaker evidence for decision making for set size 4 regardless of uniform or mixed array. Lastly, set size 4 and mixed set size 4 did not differ in P3 old-new amplitude (*t*(23) = 0.21, *p* = 0.83).

In addition to the P3 old-new amplitude, we replicated our findings in Experiment 1 and 2 that P3 latencies were significantly faster for set size 2 (435.33 ms) than for set size 4 (480.75 ms, *p* < 0.01). Different from Experiment 2, we found that mixed set size 4 had a P3 old-new latency (447.41 ms) closer to set size 2 (12.08 ms) than set size 4 (33.34 ms, *p* < 0.01). Given the role P3 old-new latency played in decision making (Hyun et al., 2008), these results suggested that the participants spent significantly less time in making decisions about a mixed set size 4 array compared to regular set size 4 array. Combining with our CDA results, we showed that participants may actively engage in category-based *removal* for mixed arrays, in that they only kept the mnenomically relevant items in working memory for comparison and decision making.

Finally, we examined whether the neural representations of visual working memory items were consistent between the retention and comparison phases. We found that CDA amplitudes in retention phase predicted those in comparison phase for set size 2 (spearman *r*(61) = 0.77, *p* < 0.001, see **Fig. 9A**) and set size 4 (spearman *r*(61) = 0.75, *p* < 0.001, see **Fig. 9B**) across individuals, suggesting that the two CDA amplitudes both reflected the number of items in visual working memory. Furthermore, we were also interested in whether participants who exhibited a larger CDA set size effect, reflected by an increase in CDA amplitude from set size 2 to set size 4, during the retention phase also showed a similar effect during the comparison phase. Pooling data from Experiments 1, 2 and 3 (n = 63), we conducted a correlational analysis to assess the relationship between the CDA increase from set size 2 to 4 during retention and comparison phases. The analysis revealed a significant positive correlation (spearman *r*(61) = 0.59, *p* < 0.001, see **Fig. 9C**), indicating that participants with a higher CDA set size effect during retention also showed a similar effect during the comparison phase. This suggests that the CDA observed during the comparison phase likely reflects the same neural signature of visual working memory as the well-established CDA during the retention phase.

**Fig 9.**
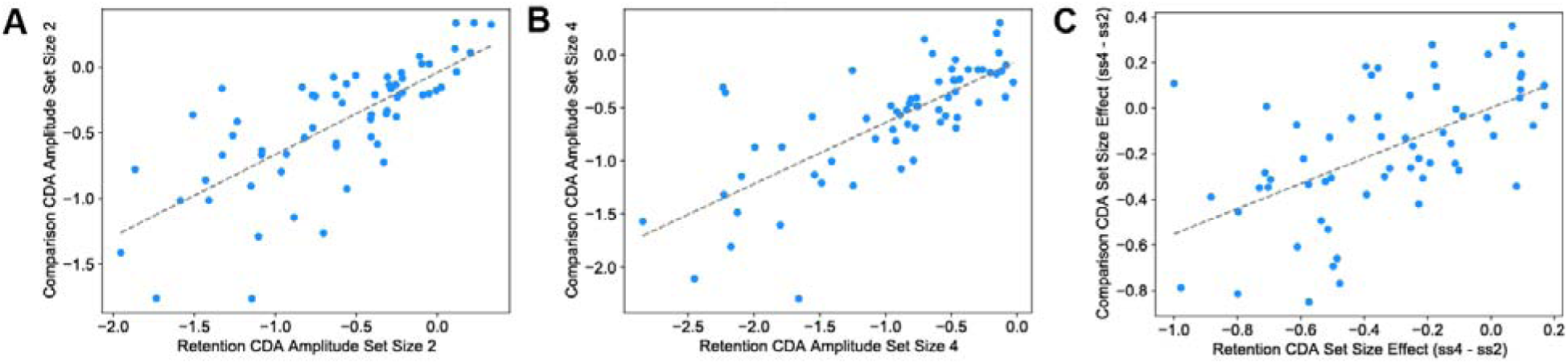
Individual Differences in CDA set size effect during retention phase predicted those during comparison phase (Exp 1-3). CDA amplitude during the retention phase predicted CDA amplitude during the comparison phase across individuals for set size 2 (Panel A), set size 4 (Panel B), and the difference between set size 4 and set size 2 (Panel C).

**Fig 10.**
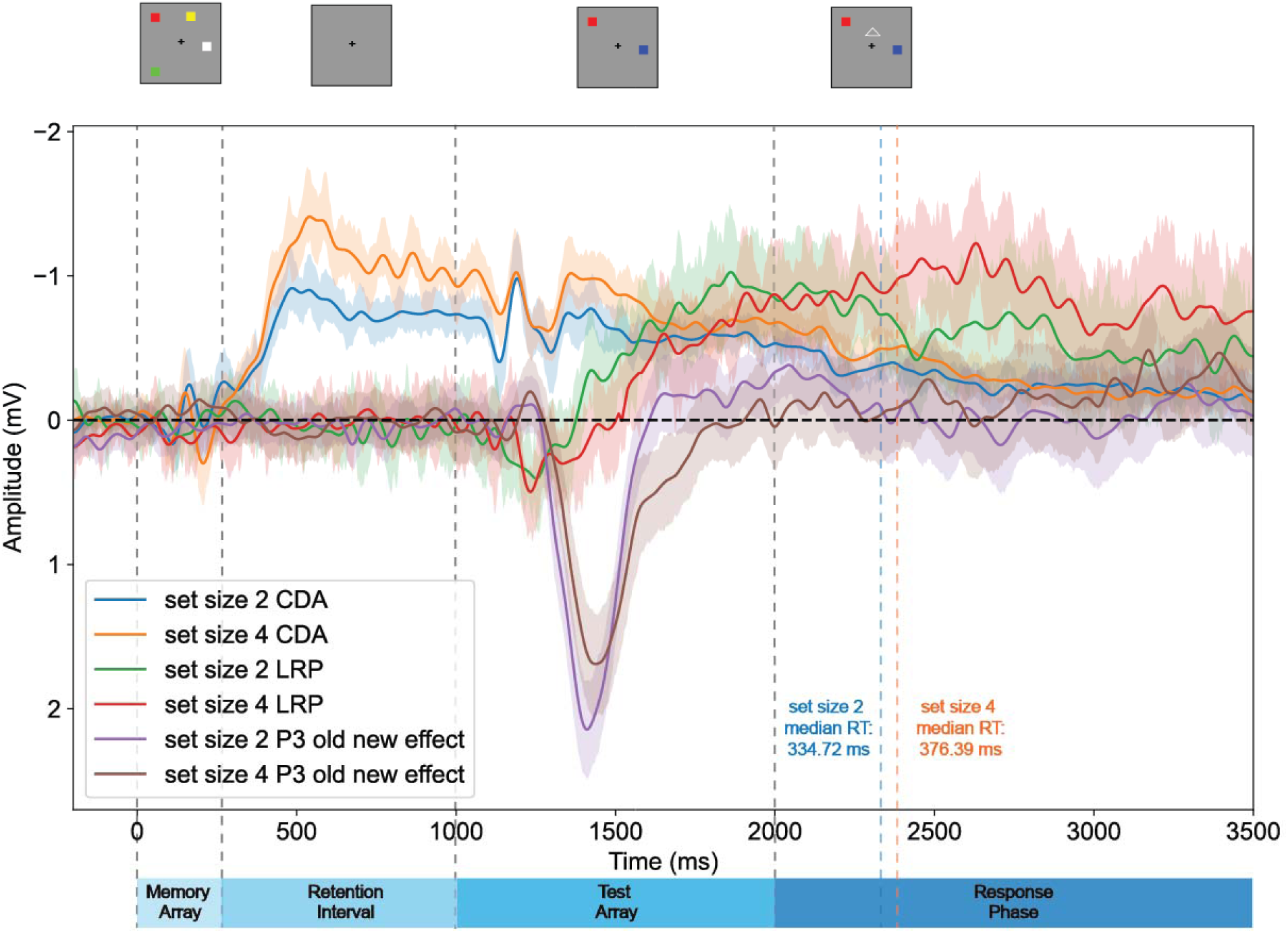
Neural working memory load (CDA) sustaind even as memory-based decision making has been made (P3 and LRP). The sustained CDA differences between set size 2 and 4 disappear shortly after a motor response has been made.

### Neural Markers of working memory at test

In our study, our primary objective was to characterize the cognitive processes underlying working memory decisions. Specifically, we examined the number of items maintained in working memory during the test phase. The contralateral delay activity (CDA), a well-established neural marker of memory load, exhibited significantly greater amplitude for set size 4 compared to set size 2 throughout the test phase (ps < 0.05 for 1000–2000 ms post-encoding array onset).

Despite this sustained memory load difference, participants completed the memory decision process, indexed by the 50% area-under-the-curve (AUC) peak latency of the P3 old-new effect, long before 2000 ms (i.e. the end of the test array with no response allowed) for both set size 2 (1419.71 ms) and set size 4 (1466.92 ms). Notably, once the decision-making process was completed, participants began preparing for a motor response, as reflected in the lateralized readiness potential (LRP), a neural index of motor preparation. In prior literature, LRP becomes more positive when a motor response is ready to be executed (Eimer 1998; Masaki et al. 2004), and had been shown to predict response time measures (Miller 1998). Our analysis revealed that LRP amplitude was significantly greater for set size 2 compared to set size 4 shortly after the disappearance of the P3 old-new effect (t(61) = 2.05, p < 0.01). These findings indicate that participants were preparing their motor responses even though CDA continued to reflect differences in memory load between set size 2 and set size 4 arrays.

Finally, after participants executed their responses (2450–3000 ms post-encoding array onset), the difference in CDA between set size 2 and set size 4 was no longer significant (*t*(61) = 0.60, p = 0.55). This suggests that working memory contents were likely discarded following response execution.

### Conclusion

In this study, we examined how people selectively remove the contents of visual working memory at test. Specifically, we asked whether participants achieve effective removal by retaining only the item relevant to the test probe or maintain the entire memory array at test. In Experiment 1, we used a single-probe change detection task to test participants’ ability to selectively access a relevant item. Despite only one item being required for comparison, CDA amplitudes scaled with set size, indicating that participants did not remove irrelevant memory contents based on spatial location, even when doing so would have been efficient for the task.

One possible explanation for the absence of removal is that maintaining the entire display may be beneficial, even when only a single probe is presented, because it preserves contextual information through relational coding (Jiang et al., 2000). In particular, retaining the original configuration in working memory may support more accurate recognition at test by allowing participants to rely on the relational structure among items. To examine the possibility that this pattern reflected spatial grouping, Experiment 2 presented two sequential memory arrays in the same spatial locations. The resulting CDA amplitudes during comparison were similar to those from a simultaneous set size 4 array, reinforcing the idea that spatial grouping alone did not guide memory removal. Consistently across Experiments 1 and 2, the P3 old-new component showed both reduced amplitude and faster latency for smaller set sizes, suggesting that decision-making was more efficient when fewer items were initially encoded, and people did not remove their memory contents at test to make decisions. Together, these findings indicate that did not remove irrelevant memory items from visual working memory based on spatial relevance at test.

Experiment 3 explored whether mnemonic category could support more selective removal at test. When participants encoded a mixed array of items from different categories (e.g., colors and shapes), CDA amplitudes during test reflected only the items that matched the category of the test probe, comparable to when only two items were originally encoded. Additionally, P3 latency was faster in these mixed-category arrays than in homogeneous arrays of the same set size, indicating more efficient access to relevant information. These results suggest that people can selectively remove certain working memory contents during retrieval based on categorical relevance, reducing memory load and enhancing decision speed. In sum, our findings show that spatial location does not guide memory removal at retrieval, but categorical information can. Rather than broadly gating out irrelevant information, participants selectively remove their working memory contents to prioritize information that is mnemonically aligned with the task demands.

Our findings suggest that participants do not spontaneously restrict retrieval in visual working memory based on spatial location when all items in an array share the same category. In these cases, rather than comparing only the item that matched the test probe’s spatial position, participants appeared to maintain all encoded items during the test phase. This lack of selective removal implies that, in the absence of competing categorical cues, participants did not remove memory contents based on spatial relevance, instead preserving the entire array for decision-making. However, when the memory array included items from distinct mnemonic categories (e.g., colors and shapes), participants retained and compared only the items matching the test probe’s category, as reflected in reduced CDA amplitudes during the comparison phase. This indicates that selective removal was guided by categorical relevance rather than spatial location.

One possibility in explaining this category-based removal is that categorical information may act as a contextual filter, helping participants efficiently isolate relevant memory representations when items from competing categories are present. This form of context-driven selection aligns with prior work on working memory access and attentional prioritization (LaRocque et al. 2013), and mirrors cue-based retrieval in long-term memory, where recognition cues preferentially trigger contextually congruent items (Schwartz et al. 2005; Polyn et al. 2009).

Another possibility is that these effects arise from encoding-based organizational strategies. When items differ categorically, participants may spontaneously structure them into separable groups, making it easier to exclude irrelevant items at test. This account is consistent with findings by Oberauer (2018), showing that removal from working memory is more effective when items are organized into a predefined subset. In our task, participants may have struggled to discard specific items from a homogeneous set (e.g., four color squares), but found it easier to limit comparison when those items were categorically distinct from the test probe.

More broadly, our results contribute to a growing body of research suggesting that mnemonic context, like category or semantic grouping, can modulate access and comparison in working memory. Prior EEG and fMRI studies have shown that items sharing the same category as a test probe can be selectively reactivated during retrieval (Rose et al. 2016), suggesting that context plays a powerful role in controlling access to memory. Our findings also offer insight into prior discrepancies in the literature. For example, earlier studies reported stable latencies in attentional components like the N2pc and posterior N2 (Talsma et al. 2001; Hyun et al. 2009) despite changes in set size. Our data suggests that while attentional selection may be engaged early on, comparison processes later in the trial depend on the number of contextually relevant items actively maintained. This interpretation is consistent with recent work showing reactivated CDA-like activity during the response phase (Roy and Faubert 2023), which likely reflects the continued maintenance of both relevant and irrelevant items unless a clear editing strategy is available.

Finally, our study helps explain why people often perform better with heterogeneous arrays in visual working memory tasks (Cai et al. 2020; Wennberg and Serences 2024). In Experiment 3, we found that the CDA amplitude during the encoding phase was similar for both homogeneous and heterogeneous arrays. This suggests that the performance benefit for heterogeneous arrays is not due to storing more information during encoding. Instead, the key difference emerged during the comparison phase: the CDA was significantly lower in the heterogeneous condition. This reduction points to a more efficient removal process, where participants may have made fewer comparison errors. One likely reason is that when items in the array are more distinct (i.e., from different categories), it’s easier to use category-based removal strategies during retrieval (Awh et al. 2007). In other words, the advantage of heterogeneity seems to come from better *selective removal* at the time of test, rather than from storing extra items in memory during encoding. Future work should explore how these *selective removal* strategies generalize across memory domains and whether they are supported by shared neural signatures in working and long-term memory (Cox and Shiffrin 2017).

## Research Transparency Statement

### General Disclosures

#### Conflicts of interest

All authors declare no conflicts of interest. Ethics: The research received approval from Institutional Review Board at The University of Chicago (IRB15-1290).

### Experiments One-Three Disclosures

#### Preregistration

No aspects of the study were preregistered. Data & Analysis scripts: The data and analysis scripts are available on Open Science Framework (https://osf.io/gd56n/).

## Acknowledgement

This research was supported by funding from the National Institute of Mental Health (Grant ROIMH087214 awarded to Edward K. Vogel), and the Office of Naval Research (Grant N00014-12-1-0972 awarded to Edward K. Vogel; Grant MURI N00014-23-1-2768 to Edward K. Vogel).

## Author Contributions

Chong Zhao played a lead role in conceptualization, data curation, formal analysis, investigation, methodology, writing–original draft, and writing–review and editing. Temilade Adekoya played a lead role in data curation and writing–review and editing. Sintra Horwitz played a lead role in data curation and writing–review and editing. Edward Awh played a lead role in conceptualization, formal analysis, investigation, methodology, funding acquisition, resources, supervision, and writing–review and editing. Edward K. Vogel played a lead role in conceptualization, formal analysis, investigation, methodology, funding acquisition, resources, supervision, and writing–review and editing.

## Notes

### Competing Interest Statement

The authors have declared no competing interest.

### Summary of Updates

We updated the manuscript extensively to incorporate the removal hypothesis throughout the introduction, results, and discussion.

