## Supplementary material for "Selective removal of visual working memory items at test": Supp.

Running title: SELECTIVE REMOVAL OF VWM AT TEST

Mailing address: 940 East 57^th^ St., Chicago, IL, 60637

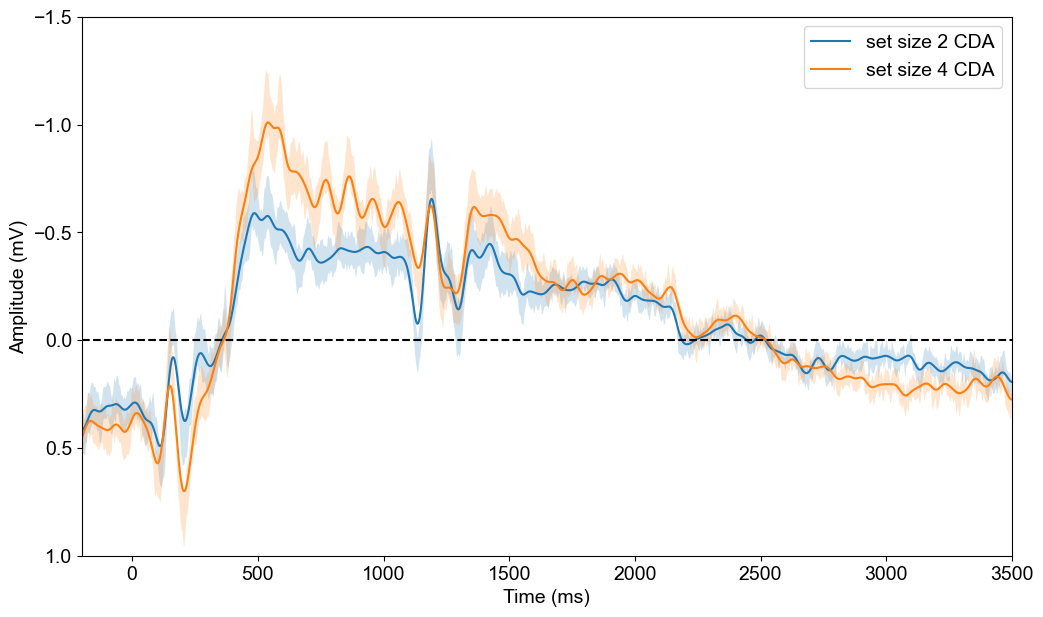

**Figure S1** Maintenance and Test CDA with post-RT baseline. Grand average ERPs showing that set-size effects remain significant through both maintenance and test phases (onset at 0ms). The CDA set size difference remains sustained past test stimulus onset (1000 ms) regardless of baseline correction.

|  | Set size 2 | Set size 4 |
| --- | --- | --- |
| Old trials | 926.06 ms | 941.13 ms |
| New trials | 889.97 ms | 989.06 ms |

**Table S1** Mean response time (ms) divided by set size and old/new in Experiment 1 for correct trials.

|  | Set size 2 | Set size 4 | Set size 4 sequential |
| --- | --- | --- | --- |
| Old trials | 637.30 ms | 711.31 ms | 660.55 ms |
| New trials | 701.07 ms | 722.21 ms | 693.10 ms |

**Table S2** Mean response time (ms) divided by set size and old/new in Experiment 2 for correct trials.

|  | Set size 2 | Set size 4 | Set size 4 mixed |
| --- | --- | --- | --- |
| Old trials | 452.99 ms | 528.83 ms | 545.38 ms |
| New trials | 493.22 ms | 572.45 ms | 530.81 ms |

**Table S3** Mean response time (ms) divided by set size and old/new in Experiment 3 for correct trials.
